# Loss of self-tolerance leads to altered gene expression and IMD pathway activation in *Drosophila melanogaster*

**DOI:** 10.1101/2021.11.19.469298

**Authors:** Pooja Kr, Ashley L. Waring-Sparks, Nicholas M. Bretz, Nathan T. Mortimer

## Abstract

Immune self-tolerance is the ability of a host’s immune system to recognize and avoid triggering immune responses against self-tissue. This allows the host to avoid self-directed immune damage while still responding appropriately to pathogen infection. A breakdown of self-tolerance can lead to an autoimmune state in which immune cells target healthy self-tissue, leading to inflammation and tissue damage. In order to better understand the basic biology of autoimmunity and the role of the innate immune system in maintaining self-tolerance, we have recently characterized the *Drosophila melanogaster tuSz* autoimmune mutant. This mutant strain can serve as a model of innate immune mediated self-tolerance, and here we identify transcripts that are deregulated in flies experiencing a loss of self-tolerance. We find that these changes include the activation of the Relish/NFκB transcription factor, alterations in transcripts encoding proteins predicted to mediate organismal metabolism, and a downregulation of transcripts linked to developmental processes. We further find that NFκB signaling plays a protective role against loss of self-tolerance in *Drosophila.* Our findings provide insight into the transcriptional and physiological changes underlying self-tolerance and autoimmunity.

## 1. INTRODUCTION

The maintenance of self-tolerance, the process in which self-tissues are identified and self-directed immune mechanisms are repressed, is an important characteristic of the immune system (Medzhitov and Janeway, 2002; Romagnani, 2006). The breakdown of the immune system’s ability to discriminate healthy self-tissue from non-self pathogens can lead to damage of a host’s own tissues by the immune system. Multiple studies have revealed the importance of self-tolerance, and the failure of this system has been implicated in a variety of autoimmune disorders and other immune mediated diseases. Although much of the research uncovering mechanisms of immune recognition and self-tolerance has focused on adaptive immune mechanisms, a growing body of work has demonstrated the importance of innate immune mechanisms in the suppression of autoimmunity (Toubi and Vadasz, 2019; Waldner, 2009).

The genetic model organism *Drosophila melanogaster* is a valuable model for understanding conserved innate immune mechanisms (Lemaitre and Hoffmann, 2007). Flies are host to a wide range of pathogens including bacteria, fungi, viruses, and parasites and use both humoral and cell mediated immune responses to combat these pathogens (Brennan and Anderson, 2004). In *Drosophila*, hemocytes (immune cells) known as plasmatocytes are found in circulation, and following stimulation, surveil the hemolymph (body cavity) for the presence of invading pathogens and tissue damage. These macrophage-like plasmatocytes are responsible for the phagocytosis of microbial pathogens and are involved in the encapsulation response against macroparasites including parasitoid wasp eggs (Honti et al., 2014; Mortimer et al., 2012; Parsons and Foley, 2016). Another immune cell type, the lamellocytes, are generally not present in a healthy fly larva, but are induced following parasitoid wasp infection and are required for melanotic encapsulation of parasitoid eggs and other foreign objects (Anderl et al., 2016; Honti et al., 2014; Mortimer et al., 2012).

*Drosophila* hemocytes also play a role in self-recognition and maintaining self-tolerance. We have recently characterized the *Drosophila tuSz^1^* temperature sensitive mutant strain which mounts a self-directed encapsulation response at the restrictive temperature (Mortimer et al., 2021). We found that the loss of self-tolerance in *tuSz^1^* is due to two genetic changes. The first is a gain of function mutation in *hopscotch* (*hop*), the *Drosophila* homolog of Janus Kinase (JAK). The JAK-STAT signaling pathway plays a key role in immune priming, and the *hop^Sz^* allele leads to hemocyte activation and the ectopic production of lamellocytes in naïve larvae (Mortimer et al., 2021). These hemocyte phenotypes are also seen in larvae with the *hop^Tum^* gain of function allele (Hanratty and Dearolf, 1993; Luo et al., 1995), but interestingly, even though *hop^Tum^* and *tuSz^1^* mutants both have hemocyte activation and excess lamellocyte production, *hop^Tum^*mutants do not display an autoimmune phenotype (Mortimer et al., 2021). This would suggest that JAK-STAT mediated immune activation alone is insufficient for the loss of self-tolerance, and that loss of self-tolerance and immune priming are genetically separable. The second mutation in the *tuSz^1^*strain is a conditional loss of function mutation in the *GCS1* gene. At the restrictive temperature, the *GCS1^Sz^* allele leads to disruption of protein N-glycosylation of extracellular matrix proteins (Mortimer et al., 2021). Together these findings define a system in which *Drosophila* self-tolerance is maintained by the ability of primed hemocytes to sense a self-signal that is encoded in the N-glycosylation of extracellular matrix proteins, and the loss of this self-recognition leads to the melanotic encapsulation of self-tissues. Thus, hemocytes can mediate both the activation of an immune response and the maintenance of self-tolerance thereby regulating immune homeostasis.

Here, we utilize the *Drosophila tuSz^1^* and *hop^Tum^*mutant strains to identify key transcriptional changes underlying immune priming, self-tolerance, and autoimmunity. Differentially expressed transcripts often reveal disease-induced changes (Porcu et al., 2021), so our analysis of transcript abundance changes in these mutants may allow us to identify processes that occur in autoimmune reactions following the loss of self-tolerance. Our results further suggest that the pro-inflammatory Rel/NFκB signaling pathway is dysregulated in the *tuSz^1^* loss of self-tolerance phenotype, and plays a protective role in this response.

## 2. MATERIALS AND METHODS

### 2.1. Insect Stocks

The *D. melanogaster* strains *w^1118^* (BDSC: 5905), *tuSz^1^* (BDSC: 5834), *PBac{GFP.FPTB-Rel}VK00037* (*Rel-GFP*) (BDSC: 81268), *hop^Tum^* (BDSC: 8492), *P{VALIUM20-GAL4.1}attP2* (*UAS-GAL4-RNAi*) (BDSC: 35784), *P{TRiP.HM05154}attP2* (*UAS-Rel-RNAi-1*) (BDSC: 28943), *P{TRiP.HMS00070}attP2* (*UAS-Rel-RNAi-2*) (BDSC: 33661), *P{TRiP.HMS00063}attP2* (*UAS-Dredd-RNAi*) (BDSC: 34070), *P{TRiP.HMC04952}attP40* (*UAS-key-RNAi*) (BDSC: 57759), and *PGRP-LB^Delta^*(BDSC: 55715) were obtained from the Bloomington Drosophila Stock Center (BDSC). *msn-GAL4* was obtained from R. Schulz (Tokusumi et al., 2009). All fly stocks were maintained on standard *Drosophila* medium on a 12-hour light-dark cycle. The study also uses the parasitoid wasp *Leptopilina boulardi* (strain Lb17) which is maintained on the *Canton S D. melanogaster* strain (BDSC: 64349).

### 2.2 RNA Isolation

Total RNA was extracted from late third instar *w^1118^*, *tuSz^1^* and *hop^Tum^* larvae raised at 28°C. In total, three biological replicates were prepared with each biological replicate consisting of 20 pooled larvae. RNA was extracted from whole larvae using Trizol (Thermo Fisher Scientific) followed by QIAGEN RNeasy Micro clean-up (QIAGEN; both according to manufacturer’s instructions). RNA concentration was measured using a nanodrop and the integrity was verified using agarose gel electrophoresis. The RNA samples were sent to Novogene (Sacramento, CA) for RNA sequencing (150bp paired end reads, Illumina). RNAseq data are available through the NCBI Gene Expression Omnibus, study GSE189383.

### 2.3 Differential Gene Expression Analysis

Sequence data were obtained in the form of FASTQ reads and analyzed using the Galaxy web platform (https://usegalaxy.org/) (Afgan et al., 2018). Initial quality control was performed using FastQC (version 0.11.8) (Andrews, 2010) to trim reads below a quality score threshold of 30. Reads were then mapped to the *D. melanogaster* reference genome dm6 using HISAT2 (version 2.1.0) (Kim et al., 2015). The resulting BAM format files were analyzed using featureCounts (version 1.6.3) (Liao et al., 2014) to generate read counts for each gene in the reference genome annotation. The counts generated by featureCounts were subsequently used to determine the differentially expressed genes using DESeq2 (version 2.11.40.6) (Love et al., 2014). The results were filtered and significant differentially expressed genes (DEGs) were identified after applying significance cut-offs of a false discovery rate (FDR) < 0.05 and log_2_ fold change ≥ 1 (for upregulated transcripts) or ≤ -1 (for downregulated transcripts).

### 2.4 Functional Annotation

The Gene Ontology (GO) Resource (http://geneontology.org/) and PANTHER classification system (Ashburner et al., 2000; Gene Ontology Consortium, 2021; Mi et al., 2013) were used for GO term analysis of each list of DEGs. The FDR threshold to determine term enrichment was 0.05.

### 2.5 Rel Localization

To determine Rel subcellular localization in hemocytes in control and mutant larvae, *w^1118^*, *tuSz^1^*, and *hop^Tum^*unmated females were crossed to *Rel-GFP* males and the crosses were raised at 28°C. From these crosses we selected *w^1118^ /Y; Rel-GFP/+* control larvae and *hop^Tum^/Y; Rel-GFP/+* and *tuSz^1^/Y; Rel-GFP/+* mutant larvae. Five male third instar larvae from each genotype were bled in PBS and hemocytes were stained using DAPI (1:500; Invitrogen) to mark nuclei and mounted in Vectashield mounting medium. To determine Rel localization in hemocytes of parasitoid infected larvae, late second instar *w^1118^ /Y; Rel-GFP/+* larvae were placed on 35mm Petri dishes filled with Drosophila medium either alone (naïve control) or together with *L. boulardi* wasps (infected) at 25°C. Larvae were dissected at 72-hours post infection (hpi; and age-matched controls) in PBS and hemocytes were stained using DAPI mounted in Vectashield for imaging. Cells were imaged using a Leica SP8 confocal microscope in the ISU Confocal Microscopy Facility to assay Rel-GFP localization.

The relative proportion of Rel-GFP signal coming from the nucleus vs cytoplasm (N/C ratio) was calculated using Fiji (Schindelin et al., 2012). Image analysis was performed as described in (Kelley and Paschal, 2019; Noursadeghi et al., 2008). Specifically, the DAPI stain was used to mask the nucleus allowing for mean cytoplasmic and nuclear fluorescence intensity values to be independently determined. The N/C ratio was then calculated based on the ratio of mean nuclear intensity to mean cytoplasmic intensity. Cells with an N/C ratio greater than 1.5 were considered to have nuclear Rel-GFP localization and cells with an N/C ratio less than 1.5 were considered to have diffuse Rel-GFP (Rel-GFP exclusion from the nucleus was not observed). This relative localization measure is a more reliable measure than absolute fluorescence intensity, being robust to stochastic cell-to-cell differences in Rel expression (Noursadeghi et al., 2008). The proportion of cells with nuclear Rel-GFP localization in the mutant or infected backgrounds was compared to *w^1118^* using Fisher’s exact test.

### 2.6 Statistical Analysis

All statistical analyses were done in the R statistical computing environment (R Core Team, 2021). The plotMDS function from the limma R package (Ritchie et al., 2015) was used to plot the first two principal components to illustrate the relationships in the expression profiles of the studied genotypes. For RNAi experiments, phenotypic penetrance data were analyzed using Fisher’s exact test, and p values were adjusted using Holm’s method to correct for multiple comparisons. Genetic interactions between *tuSz* and *PGRP-LB* were tested using ANOVA of aligned rank transformed data using the ARTool R package (Wobbrock et al., 2011) and pairwise comparisons were made using Tukey’s HSD from the multcomp R package (Hothorn et al., 2008). Graphs were produced using the ggplot2 and ggvenn R packages (Wickham, 2009; Yan, 2021) and heatmaps were produced using the gplots R package (Warnes et al., 2022).

## 3. RESULTS AND DISCUSSION

### 3.1 Identification of differentially expressed genes in *tuSz^1^* and *hop^Tum^*

To gain a better understanding of the changes involved in JAK-STAT mediated immune priming and the consequences of the loss of self-tolerance, we performed RNA sequencing on *w^1118^*, *tuSz^1^*and *hop^Tum^* larvae raised at 28°C to assess differential transcript abundance. The resulting sequence reads were mapped to the *D. melanogaster* genome (Table S1) to quantify transcript levels. We find that each genotype shows a distinct expression profile (Figure 1A), and that the genotypes are separable by plotting the first two principal components of all transcript quantification data (Figure 1B). To identify genes of interest that may be differentially expressed, we separately compared transcript levels in each mutant genotype to the *w^1118^* control genotype (Tables S2-S5). Both *tuSz^1^* and *hop^Tum^* have gain of function mutations in *hop*, so any genes that are differentially expressed in both genotypes are likely to be regulated by the JAK-STAT pathway, or to be altered because of JAK-STAT pathway-mediated changes in cell physiology. We found 832 shared genes that were differentially expressed in both *tuSz^1^* and *hop^Tum^* mutants compared to wildtype, with 384 genes that were upregulated, and 438 genes that were downregulated (Figure 1C-D). Because the JAK-STAT pathway plays a key role in immune activation (Agaisse and Perrimon, 2004), these identified genes might be relevant to the priming of the immune response.

**Figure 1.**
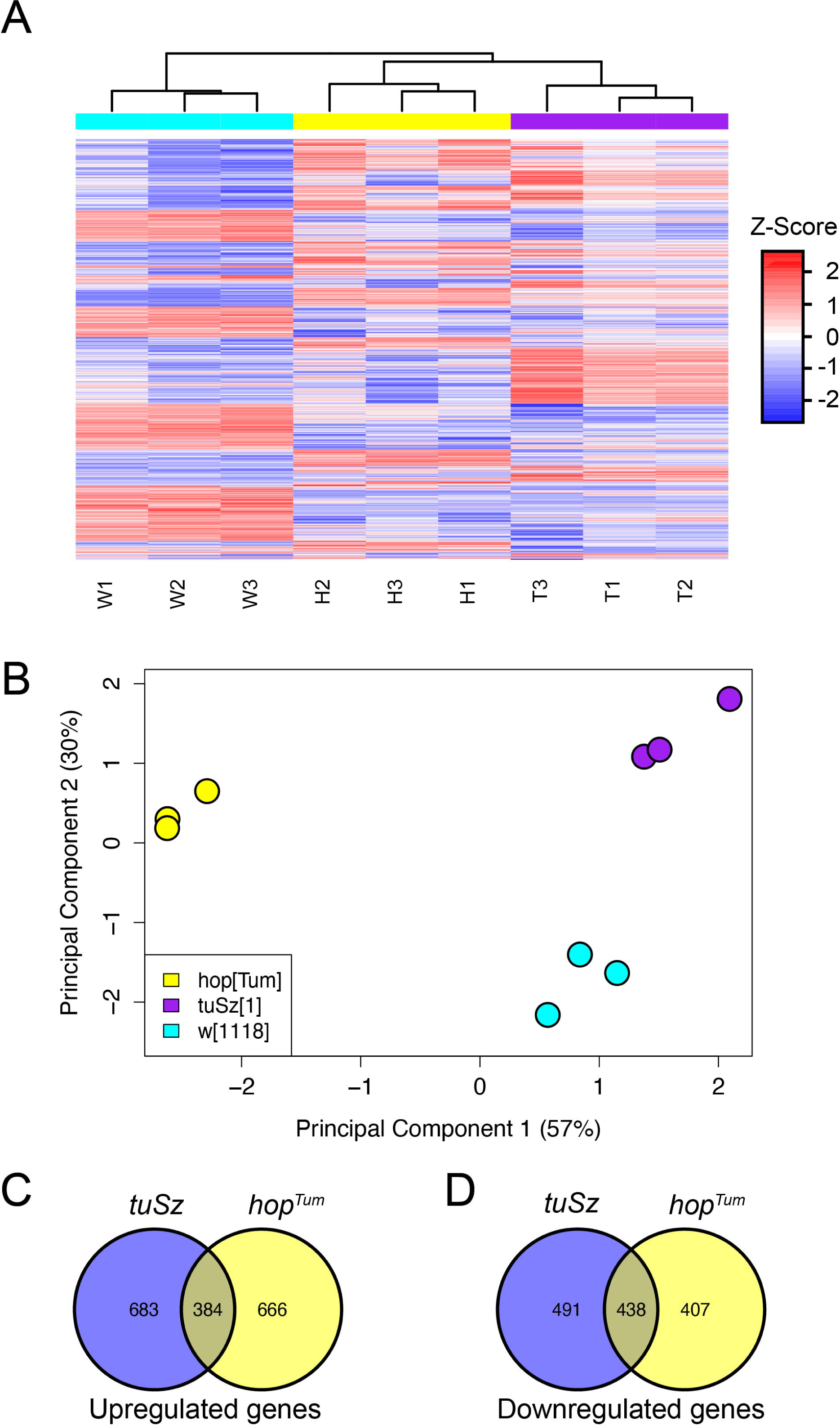
Gene expression profiles from *tuSz^1^*, *hop^Tum^* and *w^1118^*. (A) Heatmap of all genes showing altered expression between *w^1118^* and at least one mutant genotype. See Table S1 for a description of the samples. (B) Multidimensional scaling plot of the first two principal components of gene expression across the samples. (C-D) Venn diagrams illustrating the numbers of transcripts that are (C) upregulated and (D) downregulated in the indicated genotypes in comparison with *w^1118^*control larvae.

To begin characterizing these putative immune activation genes, we conducted Gene Ontology (GO) analysis to identify the biological processes and molecular functions that are enriched among the common set of differentially expressed genes (Table 1). Analysis of genes that were upregulated in both *tuSz^1^*and *hop^Tum^* identified GO terms that have been linked to host immunity including defense response to other organism (GO:0098542) and response to external stimulus (GO:0009605). General immune response genes identified in these categories (Figure 2A) include five members of the antimicrobial peptide (AMP)-like *Bomanin* gene family (*BomBc1*, *BomS1*, *BomS2*, *BomS4*, *BomT1*) (Clemmons et al., 2015) and the Cecropin family AMP gene *CecC* (Tryselius et al., 1992), along with several immune receptors (*NimB2*, *NimC1*, *NimC4*) (Kurucz et al., 2007; Somogyi et al., 2010), the prophenoloxidase family members *PPO1* and *PPO3* (Dudzic et al., 2015; Nam et al., 2008), and the *induced by infection* (*IBIN*) gene (Valanne et al., 2019). We also found differential expression of important immune related signaling pathways such as the JAK-STAT signaling pathway components *dome*, *et* and *upd3* (Myllymäki and Rämet, 2014), the Toll pathway genes *nec* and *SPE* (Kambris et al., 2006), and *Rac2* and *Rap1* which are linked to hemocyte function (Huelsmann et al., 2006; Williams et al., 2005). The *tuSz^1^* and *hop^Tum^* mutations both lead to the ectopic production of lamellocytes, and accordingly, we see lamellocyte related genes such as *ItgaPS4*, *ItgaPS5*, and *drip* upregulated in these genotypes (Stofanko et al., 2010; Tattikota et al., 2020). The proteolysis (GO:0006508) GO term is also enriched among genes upregulated in both *tuSz^1^* and *hop^Tum^*. Notably, 29 members of the S1A serine protease family are significantly upregulated in both *tuSz^1^*and *hop^Tum^*, including *CG10764* and *CG4793* which have recently been shown to be involved in the cellular immune response to parasitoid infection (Kr et al., 2021).

**Figure 2.**
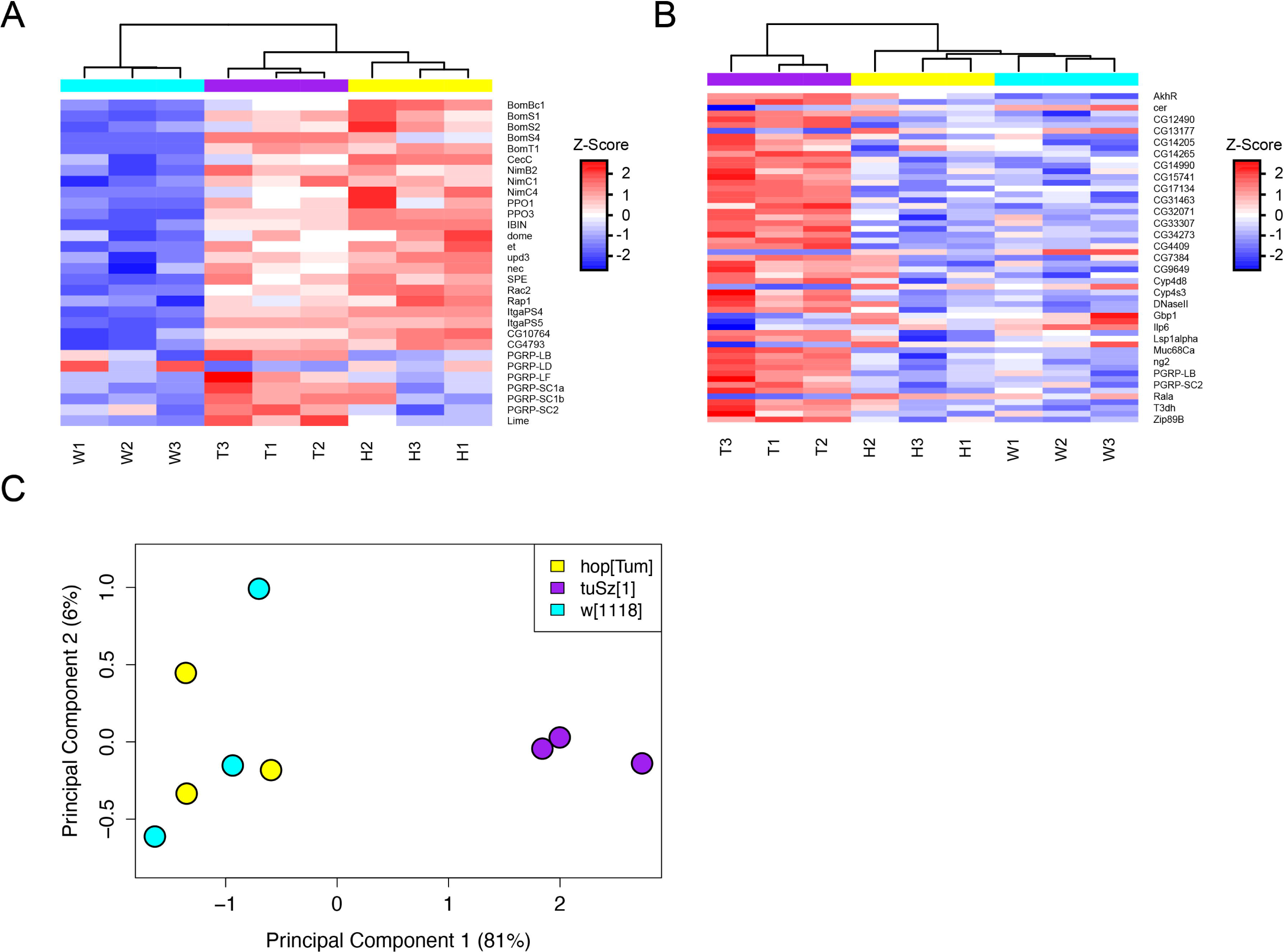
Expression of immune genes in in *tuSz^1^*, *hop^Tum^* and *w^1118^.* (A) Heatmap showing expression of representative immune genes identified by Gene Ontology analysis in *tuSz^1^*, *hop^Tum^* and *w^1118^*. See Table S1 for a description of the samples. (B) Heatmap showing expression of Imd pathway genes identified by Davoodi *et al*. (2019) in the indicated genotypes. See Table S1 for a description of the samples. (C) Multidimensional scaling plot of the first two principal components of differentially expressed Imd pathway genes between the samples.

**Table 1.** Enriched Gene Ontology (GO) Terms among DEG in common to both *tuSz^1^* and *hop^Tum^*.

| GO Term | Fold enrichment | FDR |
| --- | --- | --- |
| *Upregulated genes* |  |  |
| Defense response to other organism (GO:0098542) | 2.58 | 0.014 |
| Response to external stimulus (GO:0009605) | 2.11 | 2.6 x10<sup>-4</sup> |
| Proteolysis (GO:0006508) | 2.87 | $1.2 \times 10^{-9}$ |
| Cellular lipid metabolic process (GO:0044255) | 2.43 | 0.022 |
| *Downregulated genes* |  |  |
| Developmental process (GO:0032502) | 1.59 | $4.7 \times 10^{-4}$ |
| Regulation of multicellular organismal development (GO:2000026) | 2.24 | 0.015 |
| Tissue morphogenesis (GO:0048729) | 2.15 | 0.029 |
| Cell fate commitment (GO:0045165) | 2.99 | 0.005 |

The induction of host immunity is associated with life history tradeoffs, primarily in the redistribution of energy resources from development or reproduction to the production of an immune response (Zuk and Stoehr, 2002). We found that genes involved with cellular lipid metabolism (GO:0044255) are enriched among the common transcripts upregulated in *tuSz^1^* and *hop^Tum^*, and conversely that genes linked to development, including regulation of multicellular organismal development (GO:2000026), tissue morphogenesis (GO:0048729), and cell fate commitment (GO:0045165) are among those transcripts downregulated in both *tuSz^1^* and *hop^Tum^* (Table 1). The upregulation of transcripts involved in metabolism and downregulation of transcripts linked to developmental processes possibly reflects the immune response-development tradeoff, and suggests that this tradeoff may be regulated in part by JAK-STAT signaling or downstream processes. Further study of these genes may help to provide an understanding of the energy trade-offs and other physiological changes that occur during immune responses.

### 3.2 Loss of self-tolerance in *tuSz^1^* mutants results in differential gene expression

Despite the similarity in ectopic JAK-STAT pathway activation between the *tuSz^1^* and *hop^Tum^* mutants, *tuSz^1^*specifically displays a loss of self-tolerance phenotype (Mortimer et al., 2021). We investigated transcripts that are specifically differentially expressed in *tuSz^1^* to identify genes that might be involved in innate immune self-tolerance and autoimmune mechanisms. Out of 1174 transcripts that were differentially expressed in *tuSz^1^*but not *hop^Tum^* (referred to as *tuSz^1^* specific DEGs), we find 683 transcripts that are significantly upregulated and 491 transcripts that are significantly downregulated (Figure 1C-D). It is likely that this class of *tuSz^1^* specific DEGs represents genes that are altered due to the physiological changes seen in larvae during the autoimmune response to loss of self-tolerance rather than direct targets of JAK-STAT signaling.

In contrast to the list of upregulated transcripts in common to *tuSz^1^* and *hop^Tum^*, GO Term analysis of these *tuSz^1^* specific DEGs (Table 2) failed to find enrichment of any GO categories associated with positive regulation of host immunity. Indeed, one of the most highly enriched terms was the negative regulation of antimicrobial peptide biosynthetic process (GO:0002806). This term was linked to peptidoglycan recognition protein (PGRP) encoding genes, and we find that six PGRP gene transcripts are altered in *tuSz^1^*, but not *hop^Tum^*, mutants (Figure 2A). There are 13 members of this family in the *Drosophila melanogaster* genome (Table 3), which primarily play roles in the regulation of the Toll and Imd immune signaling pathways (Kurata, 2014). There are five PGRP genes that encode negative regulators of Imd signaling, and we find that all five of these are specifically upregulated in *tuSz^1^* mutants, with the sixth altered PGRP playing an unknown role (Table 3). The finding that whereas positive regulators of immunity are identified as upregulated genes in both *tuSz^1^* and *hop^Tum^*, the enrichment of genes involved in negative regulation of the Imd pathway specifically in *tuSz^1^*mutants suggests a differential role of immune mechanisms in JAK-STAT mediated immune priming and in the autoimmune response to a loss of self-tolerance. More specifically, the regulatory PGRP genes we find elevated specifically in *tuSz^1^* larvae are induced downstream of Imd signaling in a negative feedback regulatory loop (Zaidman-Rémy et al., 2006). This finding suggests that Imd pathway activity may be altered in the autoimmune environment resulting from the loss of self-tolerance in *tuSz^1^*larvae.

**Table 2.** Enriched GO Terms among *tuSz^1^* specific DEGs.

| GO Term | Fold enrichment | FDR |
| --- | --- | --- |
| *Upregulated genes* |  |  |
| Negative regulation of antimicrobial peptide biosynthetic process (GO:0002806) | 12.42 | 0.031 |
| Chitin-based cuticle development (GO: 0040003) | 2.89 | $4.9 \times 10^{-6}$ |
| Oxidoreductase activity (GO: 0016491) | 2.80 | $5.4 \times 10^{-13}$ |
| Generation of precursor metabolites and energy (GO:0006091) | 2.32 | 0.023 |
| Carbohydrate derivative metabolic process (GO:1901135) | 2.14 | $5.0 \times 10^{-5}$ |
| Alpha-amino acid metabolic process (GO:1901605) | 3.29 | 0.006 |
| Lipid metabolic process (GO:0006629) | 1.98 | 0.003 |
| *Downregulated genes* |  |  |
| Tissue morphogenesis (GO:0048729) | 2.43 | $2.3 \times 10^{-4}$ |
| Establishment of tissue polarity (GO:0007164) | 4.17 | 0.049 |
| Appendage development (GO:0048736) | 2.63 | 0.004 |
| Central nervous system development (GO:0007417) | 2.53 | 0.042 |
| Cell differentiation (GO:0030154) | 1.53 | 0.043 |
| Pattern specification process (GO:0007389) | 2.09 | 0.035 |
| Regulation of cell communication (GO:0010646) | 1.83 | 0.025 |
| Positive regulation of stem cell proliferation (GO:2000648) | 9.6 | 0.003 |
| Metamorphosis (GO:0007552) | 2.42 | 0.002 |

**Table 3.**
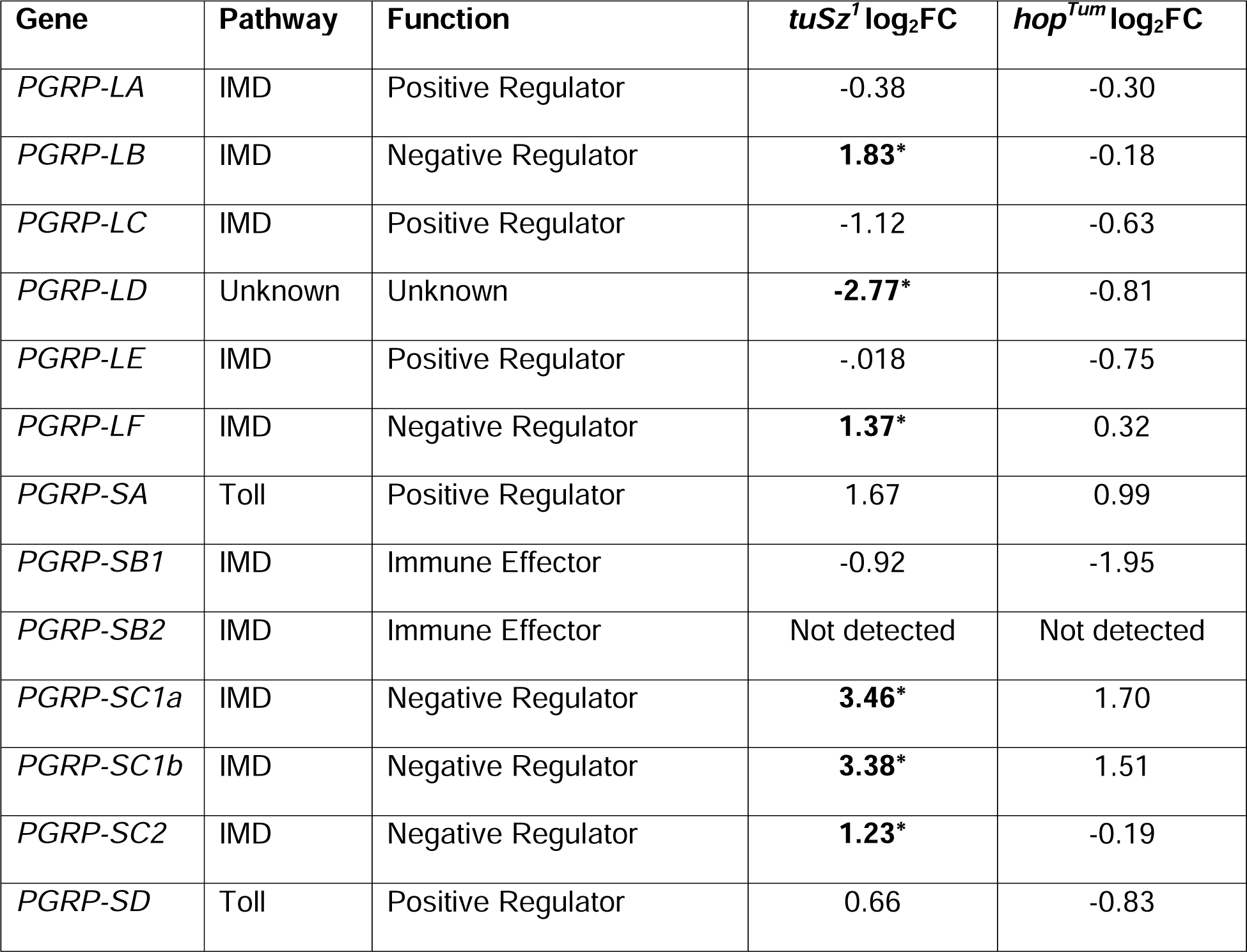
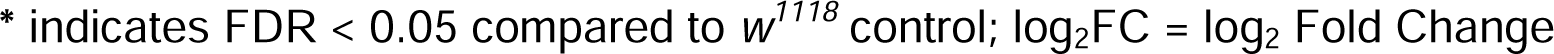
Expression of PGRP family genes in *tuSz^1^* and *hop^Tum^* compared to *w^1118^* control larvae.

| Gene | Pathway | Function | *tuSz<sup>1</sup>* log<sub>2</sub>FC | *hop<sup>Tum</sup>* log<sub>2</sub>FC |
| --- | --- | --- | --- | --- |
| *PGRP-LA* | IMD | Positive Regulator | -0.38 | -0.30 |
| *PGRP-LB* | IMD | Negative Regulator | **1.83\*** | -0.18 |
| *PGRP-LC* | IMD | Positive Regulator | -1.12 | -0.63 |
| *PGRP-LD* | Unknown | Unknown | **-2.77\*** | -0.81 |
| *PGRP-LE* | IMD | Positive Regulator | -0.018 | -0.75 |
| *PGRP-LF* | IMD | Negative Regulator | **1.37\*** | 0.32 |
| *PGRP-SA* | Toll | Positive Regulator | 1.67 | 0.99 |
| *PGRP-SB1* | IMD | Immune Effector | -0.92 | -1.95 |
| *PGRP-SB2* | IMD | Immune Effector | Not detected | Not detected |
| *PGRP-SC1a* | IMD | Negative Regulator | **3.46\*** | 1.70 |
| *PGRP-SC1b* | IMD | Negative Regulator | **3.38\*** | 1.51 |
| *PGRP-SC2* | IMD | Negative Regulator | **1.23\*** | -0.19 |
| *PGRP-SD* | Toll | Positive Regulator | 0.66 | -0.83 |

\* indicates FDR < 0.05 compared to $w^{1118}$ control; log<sub>2</sub>FC = log<sub>2</sub> Fold Change

We additionally find 26 genes that map to the chitin-based cuticle development (GO: 0040003) term among the *tuSz^1^* specific DEGs, including members of the *Lcp*, *Acp*, *Twdl*, *Cpr*, and *Ccp* gene families (Table 2). Expression of cuticle genes has been linked to infection in *D. melanogaster* and other insects, although a role in host immunity has yet to be identified (Hou et al., 2021; Pantha et al., 2021; Yadav et al., 2017). Many of the additional enriched GO categories among the *tuSz^1^* specific DEGs are associated with stress responses and metabolism, perhaps providing insight into physiological changes in *tuSz^1^* mutant larvae due to the occurrence of an autoimmune response (Table 2). We find an enrichment of oxidoreductase activity (GO: 0016491) genes, including the glutathione transferase (GST) encoding genes *GstD2, GstD6, GstD7, GstD8, GstE1, GstE3, GstE5,* and *GstE7* (Low et al., 2007). We also find that 18 cytochrome P450 (CYP450) encoding genes are upregulated in *tuSz^1^* mutants. CYP450s are involved in a wide range of cellular activities including roles in hormone signaling and detoxification of xenobiotics (Bergé et al., 1998; Chung et al., 2009). The specific upregulation of these stress response genes suggests that loss of self-tolerance may lead to the production of an internal oxidizing environment.

As mentioned previously, metabolic changes often accompany immune responses, and we find that two general regulators of host metabolism, the *Adipokinetic hormone receptor* (*AkhR*) (Bharucha et al., 2008) and *Linking immunity and metabolism* (*Lime*) (Figure 2) (Mihajlovic et al., 2019), are upregulated in *tuSz^1^* mutants. *Lime* encodes a transcription factor that plays a key role in metabolic changes following infection. Accordingly, we find that genes associated with a wide range of metabolic activities including generation of precursor metabolites and energy (GO:0006091), carbohydrate derivative metabolic process (GO:1901135), alpha-amino acid metabolic process (GO:1901605), and lipid metabolic process (GO:0006629) were enriched among those upregulated in *tuSz^1^* larvae (Table 2). Further analysis of these genes and the mechanisms they mediate will help to identify the metabolic pathways involved in the response to a loss of self-tolerance.

Along with upregulated transcripts, transcripts that are downregulated in these genotypes may also provide insight into the cellular activities that are altered. We find that genes associated with tissue morphogenesis (GO:0048729) and the establishment of tissue polarity (GO:0007164) are enriched among the downregulated *tuSz^1^* specific DEGs (Table 2). The loss of cellular polarity and disrupted tissue organization has also been observed in both autoimmune mouse models and patients with autoimmune conditions such as inflammatory bowel disease (Ahmad et al., 2017; Guo and Shen, 2021; Klunder et al., 2017). This may suggest that tissue structural maintenance is a key self-tolerance checkpoint, in agreement with previous findings in *Drosophila* (Kim and Choe, 2014; Rizki and Rizki, 1974).

### 3.3 Relish/NF**_κ_**B is activated in immune tissues in *tuSz^1^* mutants

Based on the specific upregulation of Imd-responsive PGRP genes (Table 3), we hypothesized that Imd signaling may be dysregulated in the *tuSz^1^* loss of self-tolerance phenotype. While the role of the Imd pathway in mediating the *Drosophila* humoral immune response to Gram-negative bacteria has been well established (Lemaitre et al., 1995), the mechanism by which it may participate in the response to loss of self-tolerance is unknown. The *tuSz^1^* autoimmune mutant may therefore provide an effective model to begin exploring the role of the Imd pathway in self-tolerance and autoimmune reactions.

The Imd pathway mainly responds to infection via differential expression of target genes (Kleino and Silverman, 2014). A previous study has identified genes whose expression is modified by constitutive Imd pathway activation (Davoodi et al., 2019). We find that 58 of the 324 genes identified in this study also showed differential expression in *tuSz^1^*larvae (Figure 2B) (Table S6), including three of the identified PGRP genes (*PGRP-LB*, *PGRP-LF*, *PGRP-SC2*), and other immune genes such as *AttA, Rala, Gbp1* and *GILT2*. The altered expression of Imd pathway genes is not seen in *hop^Tum^*mutants (Figure 2B), and the *tuSz^1^* expression profile is clearly distinct from both *w^1118^* and *hop^Tum^* by principal components analysis of Imd pathway transcript levels (Figure 2C). Notably, *w^1118^*and *hop^Tum^* samples are indistinguishable in this analysis, suggesting minimal alteration of Imd signaling in *hop^Tum^*. These findings provide support for the hypothesis that Imd signaling is dysregulated in *tuSz^1^* mutants in response to the loss of self-tolerance rather than as a direct result of JAK-STAT pathway activation. The overlap of gene expression changes between *tuSz^1^* mutants and constitutive Imd activation suggests that the Imd pathway is activated in *tuSz^1^* mutant larvae.

Gene expression downstream of Imd is largely mediated by the activation of Relish (Rel), a *D. melanogaster* member of the NF-κB family of transcriptional activators (Hetru and Hoffmann, 2009). Rel activation is regulated by signal-dependent translocation into the nucleus. Whereas full-length Rel protein is autoinhibited and remains in the cytoplasm, Imd activation results in cleavage of Rel and the production of an active N-terminal fragment, which translocates to the nucleus to drive gene expression (Stöven et al., 2000). To determine whether Rel is activated in *tuSz^1^*mutants, we used a *Rel-GFP* fusion protein encoding transgenic line to track the subcellular localization within the main immune tissues, the hemocytes and fat body. This reporter has GFP fused to the N-terminal domain of Relish, and GFP localization in the nucleus versus cytoplasm gives a relative readout of pathway activity.

*Rel-GFP* expression was observed in plasmatocytes in *tuSz^1^/Y; Rel-GFP/+* and *w^1118^ /Y; Rel-GFP/+* larvae (Figure 3A-B). We found that *Rel-GFP* was significantly more likely to be concentrated in the nucleus in plasmatocytes from *tuSz^1^/Y; Rel-GFP/+* larvae compared to *w^1118^ /Y; Rel-GFP/+* control larvae (Fisher’s exact test: p = 1.2x10^-5^; odds ratio = 9.02) or compared to *hop^Tum^* mutant larvae (Fisher’s exact test: p = 9.7x10^-4^; odds ratio = 5.87) (Figure 3D), suggesting that *Rel* activation is induced specifically in *tuSz^1^*mutants. In addition, *Rel-GFP* expression and nuclear localization were observed in lamellocytes from *tuSz^1^/Y; Rel-GFP/+* larvae (Figure 3C), suggesting that the Imd pathway is also active in this hemocyte type in the *tuSz^1^*mutant background.

**Figure 3.**
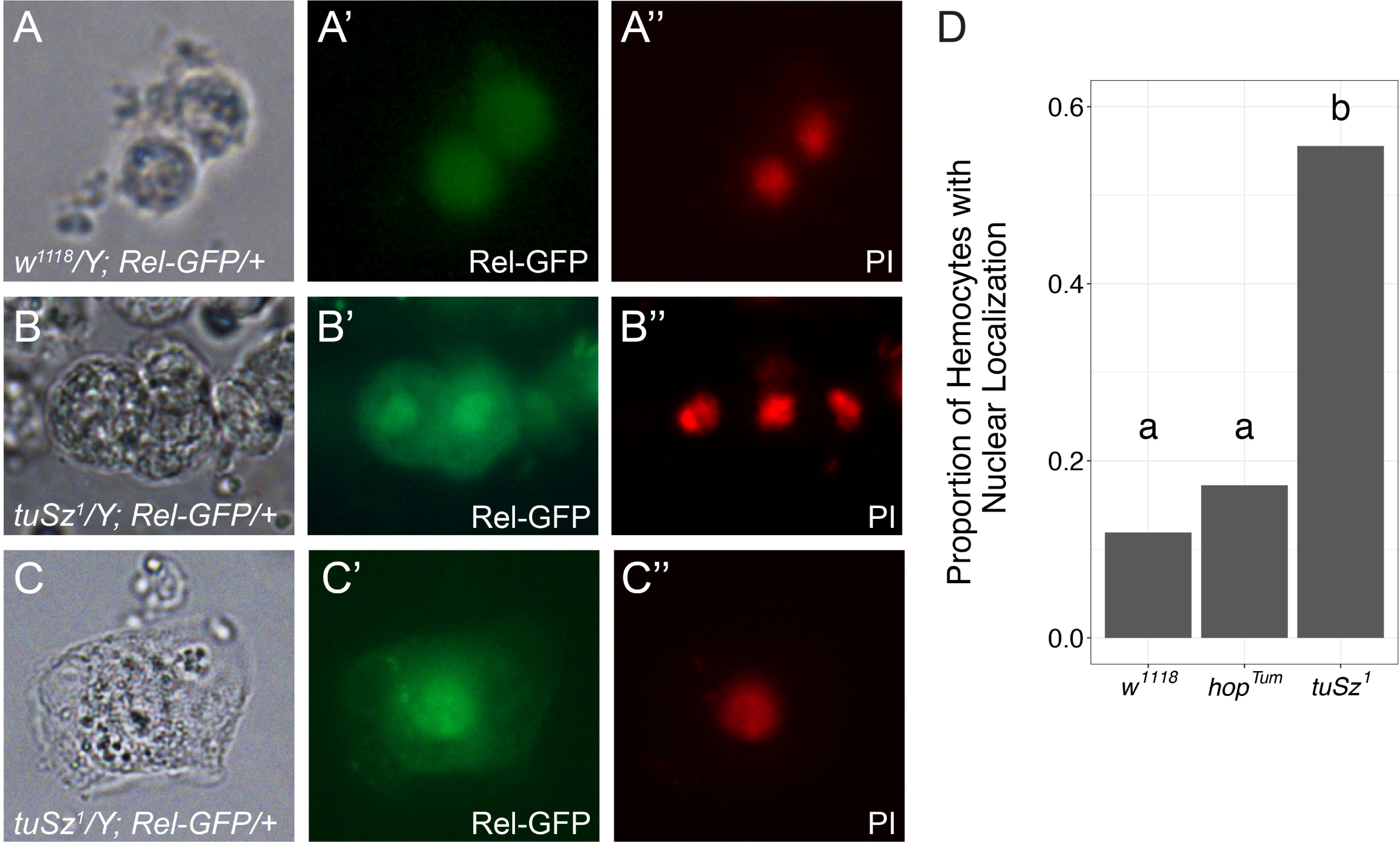
Localization of the Rel-GFP fusion protein in hemocytes. Hemocytes were dissected from *w^1118^/Y*; *Rel-GFP/+* and *tuSz^1^/Y*; *Rel-GFP/+* larvae raised at 28°C, fixed, and stained with propidium iodide (PI). Shown are representative images of hemocytes: brightfield (A-C), GFP to visualize Rel-GFP localization (A’-C’) and PI to mark nuclei (A’’-C’’). Plasmatocytes are shown in A and B, lamellocytes are shown in C. (D) Bar graph showing the proportion of hemocytes with a predominantly nuclear Rel-GFP localization in the indicated genotypes (see Materials and Methods for analysis details). Complete genotypes *tuSz^1^/Y*; *Rel-GFP/+, hop^Tum^/Y; Rel-GFP/+* and *w^1118^/Y*; *Rel-GFP/+* are simplified to *tuSz^1^*, *hop^Tum^*, and *w^1118^* for clarity. Distinct letter labels denote differences with p < 0.05.

To determine if this Rel activation is common to cellular immune responses in *Drosophila* or if it may be specific to loss of self-tolerance reactions, we assayed *Rel-GFP* localization in the immune cells of *w^1118^ /Y; Rel-GFP/+* larvae during the response to parasitoid wasp infection. *w^1118^ /Y; Rel-GFP/+* larvae were infected by *Leptopilina boulardi* and hemocytes were isolated and imaged 72hpi. We observed minimal *Rel-GFP* expression in both plasmatocytes (Figure 4A-B) and lamellocytes (Figure 4C) in these infected larvae. Circulating non-hemocyte cells (marked by an asterisk* in Figure 4B-C) showed strong fluorescence and acted as an imaging control. These results suggest that Rel and the Imd pathway are not likely to be involved in the cellular immune response to parasitoid infection. In agreement with this finding, a previous study has shown that *Rel* mutants have a normal encapsulation response against parasitoid wasp infection (Hedengren et al., 1999).

**Figure 4.**
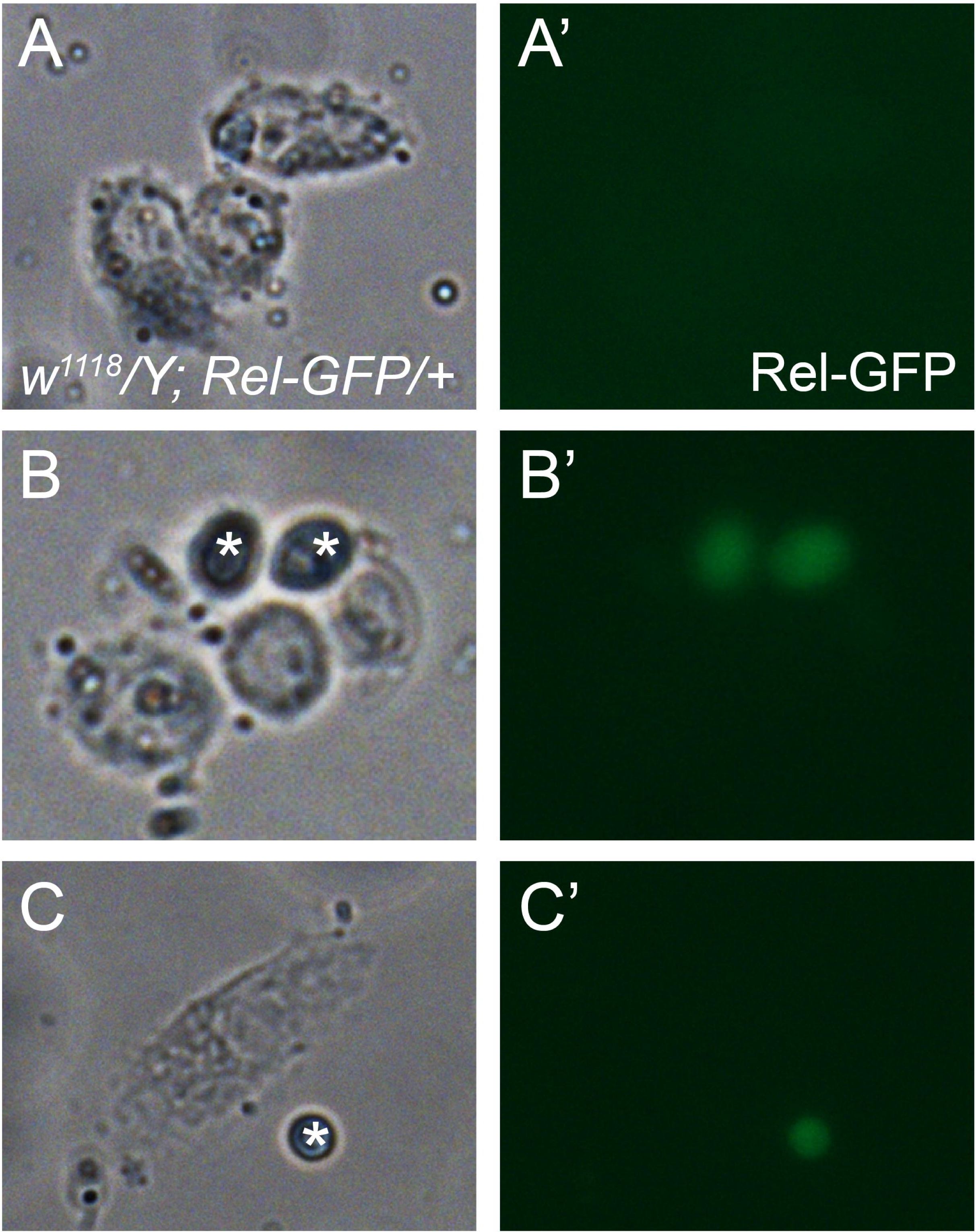
Expression of the Rel-GFP fusion protein in hemocytes from parasitoid infected *w^1118^/Y*; *Rel-GFP/+* larvae. Brightfield (A-C) and GFP (A’-C’) images. Rel-GFP expression in an unknown non-hemocyte cell type is indicated by * (B-C).

To test whether loss of *Rel* modifies the *tuSz^1^*loss of self-tolerance phenotype, we used RNA interference (RNAi) to knock-down *Rel* transcripts in immune cells in the *tuSz^1^* mutant background. In comparison with a genetic control (*GAL4-RNAi*), the expression of either of two distinct *Rel* knockdown constructs (*Rel-RNAi-1* and *Rel-RNAi-2*) in immune cells leads to an increased penetrance of the autoimmune phenotype (*Rel-RNAi-1*: Fisher’s exact test: p = 4.3x10^-3^; *Rel-RNAi-2*: Fisher’s exact test: p = 3.7x10^-5^) (Figure 5A). To further test the role of *Rel* activation, we used knockdown constructs targeting two distinct positive regulators of *Rel*: *key,* a component of the IκB Kinase complex which is required for *Rel* activation (Ertürk-Hasdemir et al., 2009) (*key-RNAi*) and *Dredd,* the caspase that directly cleaves and activates *Rel* (Stöven et al., 2003) (*Dredd-RNAi*). Similar to *Rel-RNAi-1* and *-2, Dredd-RNAi* mediated knockdown enhances the penetrance of the *tuSz^1^* phenotype (Fisher’s exact test: p = 1.0x10^-6^) (Figure 5A). Additionally, *key-RNAi* mediated knockdown shows considerable lethality in the *tuSz^1^* background, which we have previously observed with mutations that strongly enhance the *tuSz^1^*phenotype (Mortimer et al., 2021). These results suggest that *Rel*/NFκB transcriptional activity is protective against loss of self-tolerance.

**Figure 5.**
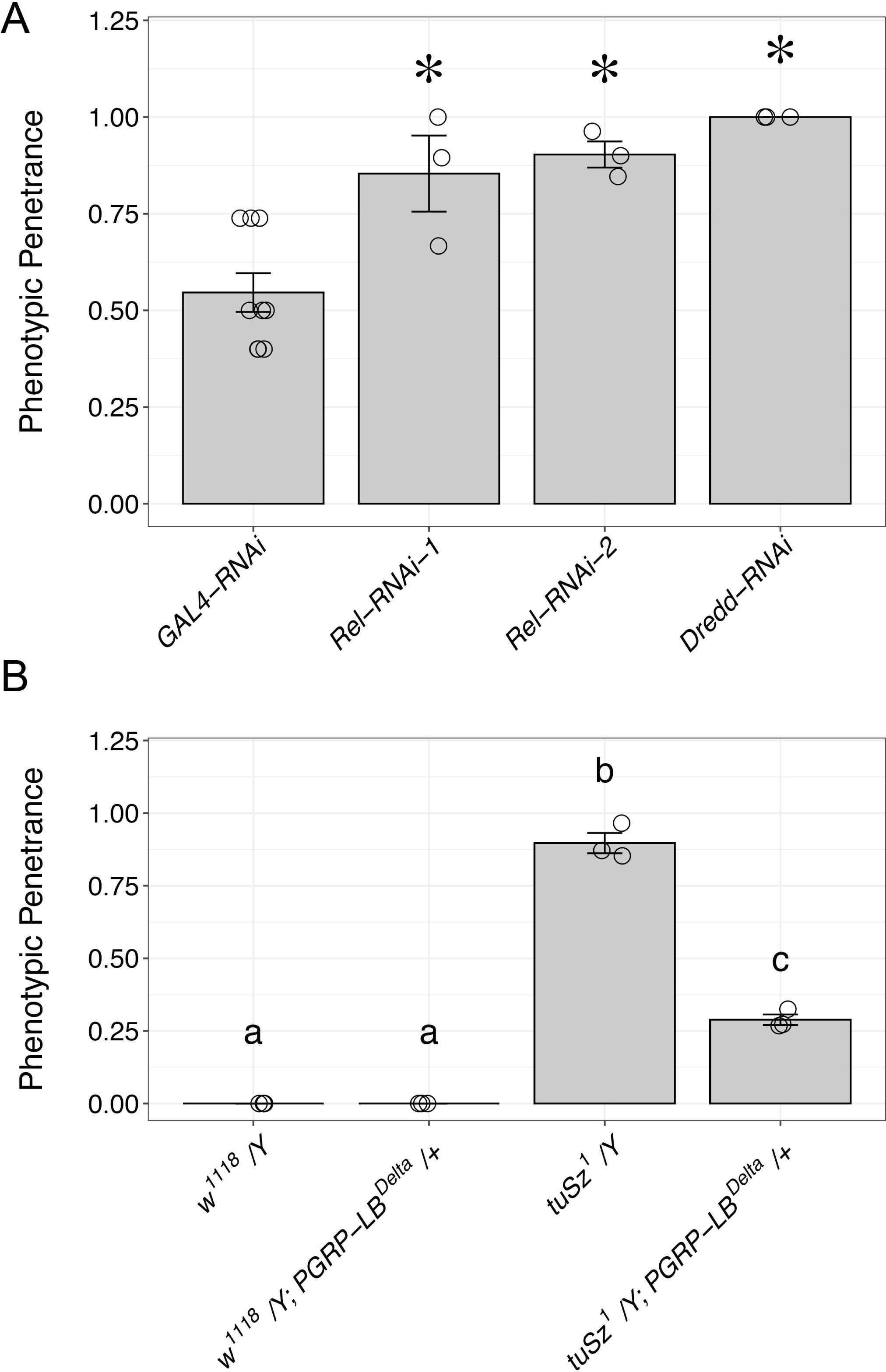
*Rel/*NFκB signaling modifies the *tuSz^1^* phenotype. (A) Penetrance of the *tuSz^1^* phenotype in adult flies of the indicated genotypes. Data are plotted as the average penetrance ± standard error across performed replicates. The mean penetrance of each replicate is visualized with open circles. * p < 0.05 compared to *GAL4-RNAi*. (B) Penetrance of the *tuSz^1^* phenotype in adult flies of the indicated genotypes. Data are plotted as the average penetrance ± standard error across performed replicates. The mean penetrance of each replicate is visualized with open circles. Distinct letter labels denote differences with p < 0.05 by Tukey’s HSD.

*Rel*/NFκB transcriptional activity is associated with pro-inflammatory signaling in *Drosophila* and other species (Ganesan et al., 2011; Lawrence, 2009; Libert et al., 2006; Myllymäki et al., 2014). The observed Rel activation suggests that inflammation is a feature of loss of self-tolerance responses in *Drosophila*, a mechanism that is conserved in human autoimmune disease (Furman et al., 2019; Zhang et al., 2017). The finding that loss of Rel activation enhances the *tuSz^1^*phenotype is surprising given that the constitutive activation of NF-κB is implicated in the pathogenesis of multiple forms of autoimmune disease (Ilchovska and Barrow, 2021; Miraghazadeh and Cook, 2018; Sun, 2017; Zhang et al., 2017). However, loss of mutations in NF-κB family genes including the *Rel* homolog RELA have also been linked to the development of autoimmune disorders (Barnabei et al., 2021; Comrie et al., 2018). This suggests that, in some situations, NF-κB signaling can be protective against disruptions in self-tolerance and progression of autoimmune disorders. Our findings of the protective effect of *Rel* in *Drosophila* may provide a model to further explore this underappreciated role of NFκB signaling.

To better understand the role of Rel activation in the response to loss of self-tolerance in *Drosophila*, we tested the effect of a loss of function mutation in the negative regulatory PRGP family protein PGRP-LB. We find that in the *tuSz^1^* genetic background, a heterozygous deletion mutation of the *PGRP-LB* gene dominantly and significantly suppresses the *tuSz^1^* mutant phenotype (Full genotype: *tuSz^1^/Y*; *PGRP-LB^Delta^/+*. Tukey’s HSD: t = -5.196; p = 3.7x10^-3^) (Figure 5B). Interestingly, the loss of one copy of the *PGRP-LB* gene in an otherwise wildtype background does not lead to a visible phenotype (Full genotype: *w^1118^/Y*; *PGRP-LB^Delta^/+*. Tukey’s HSD: t = 0.0; p = 1.0) (Figure 5B).

The negative regulatory PGRPs are amidase or amidase-like proteins that suppress the Imd/*Rel* pathway by either degrading (amidase PGRPs) or scavenging (amidase-like PGRPs) peptidoglycans (PGNs), the bacterial cell well components that trigger pathway activation (Bischoff et al., 2006; Maillet et al., 2008; Zaidman-Rémy et al., 2006). PGRP-LB is an active amidase that is specific for *meso-*diaminopimelic acid-containing peptidoglycans (DAP-PGN), which are found predominantly in gram negative bacteria (Zaidman-Rémy et al., 2006). Based on this mode of action, we would not predict that loss of PGRP-LB would have an impact in the sterile inflammation environment of *tuSz^1^*mutants. However, structural analysis of PGRPs suggests a mechanism through which these proteins may have target-binding specificity beyond the PGN sub-type (Kim et al., 2003; Royet et al., 2011). Multiple examples of PGN-independent activity have also been observed in diverse species (Lu et al., 2006; Sharma et al., 2011; Tydell et al., 2006). The loss of self phenotype in *tuSz^1^* mutants arises due to defects in the N-glycosylation of extracellular matrix proteins (Mortimer et al., 2021). Perhaps these mis-glycosylated proteins serve as an altered-self signal (Maverakis et al., 2015), which is recognized by PGRPs as an additional, and thus far unidentified, target. If PGRP-LB recognizes mis-glycolysated proteins to trigger an immune response, it could explain the strong suppression of the tuSz^1^ mutants by heterozygosity for the *PGRP-LB* gene. Conversely, PGRPs have also been linked to modulating host immunity and tolerance to the gut microbiota (Bosco-Drayon et al., 2012). It may be that the loss of *PGRP-LB* leads to alterations in the gut microbiome, which may have direct or indirect effects on the loss of self-tolerance immune response that characterizes *tuSz^1^*mutants.

## 4. CONCLUSION

Immune self-tolerance is essential for maintaining immune homeostasis, and breakdown of self-tolerance mechanisms results in autoimmunity and other immune related diseases. Although great progress has been made in understanding self-tolerance in the context of adaptive immunity, innate immune mechanisms regulating self-tolerance and its role in autoimmunity remain less well characterized. In this study we have identified genes that are differentially expressed following loss of self-tolerance in the *Drosophila melanogaster tuSz^1^* mutant. These results provide insight into the biological processes that take place in *tuSz^1^* mutant larvae and may contribute toward a better understanding of the physiological changes in loss of self-tolerance autoimmune reactions.

Our findings also serve to highlight the complex relationship between the regulation of immune signaling mechanisms and the development of autoimmune phenotypes. We find that, similar to human autoimmune disease, the loss of self-tolerance in *Drosophila* leads to elevated *Rel*/NFκB signaling. Dysregulation of NFκB through both ectopic activation or loss of function is associated with the pathogenesis of autoimmunity, and we find that the pathway plays a protective role in *Drosophila*. This will provide a tractable genetic model for further studies into the complex roles of NFκB in autoimmune disorders. Additionally, we have identified a role for a peptidoglycan recognition protein in regulating the response to a loss of self-tolerance in the absence of bacterial peptidoglycans. These PGRPs are widely conserved from insects to vertebrates, but their possible roles outside of bacterial pattern recognition have yet to be explored. We have identified the intriguing possibility that certain PGRP proteins may recognize self or lack-of-self epitopes and contribute to loss of self-tolerance reactions. Again, the loss of self-tolerance *Drosophila* mutant lines may provide a model to better understand these functions.

## ACKNOWLEDGEMENTS

We thank Joshua Hill and Elise N. Le for their assistance on this project. The ISU Confocal Microscopy Facility was funded by NSF grant DBI-1828136. Stocks obtained from the Bloomington Drosophila Stock Center (NIH P40OD018537) were used in this study.

