## Supplementary material for "Loss of self-tolerance leads to altered gene expression and IMD pathway activation in *Drosophila melanogaster*": Table S1

**Table S1**. RNAseq summary with the total number of reads and the number and proportion of reads mapped to the *Drosophila melanogaster* genome from each replicate sample.

| **Genotype (Replicate)** | **Number of Reads** | **Number of Reads Mapped to the *D. melanogaster* genome** | **Percentage of Reads Mapped to the *D. melanogaster* genome** |
| --- | --- | --- | --- |
| *tuSz^1^* (T1) | 21711455 | 17710847 | 81.6% |
| *tuSz^1^* (T2) | 23884488 | 20406116 | 85.4% |
| *tuSz^1^* (T3) | 26169935 | 21815202 | 83.4% |
| *hop^Tum^* (H1) | 23040246 | 18995520 | 82.4% |
| *hop^Tum^* (H2) | 26165030 | 20867356 | 79.8% |
| *hop^Tum^* (H3) | 22702836 | 18450982 | 81.3% |
| *w^1118^* (W1) | 23692195 | 19860760 | 83.8% |
| *w^1118^* (W2) | 22490319 | 19392201 | 86.2% |
| *w^1118^* (W3) | 23950610 | 18867852 | 78.8% |
